# GATA2 haploinsufficiency causes an epigenetic feedback mechanism resulting in myeloid and erythroid dysplasi

**DOI:** 10.1101/2021.10.29.466416

**Authors:** Emanuele Gioacchino, Wei Zhang, Cansu Koyunlar, Joke Zink, Hans de Looper, Kirsten J. Gussinklo, Remco Hoogenboezem, Dennis Bosch, Eric Bindels, Ivo P. Touw, Emma de Pater

## Abstract

The transcription factor GATA2 has pivotal roles in hematopoiesis. Germline GATA2 mutations in patients result in GATA2 haploinsufficiency syndrome characterized by immunodeficiency, bone marrow failure, and predispositions to myelodysplastic syndrome (MDS) and acute myeloid leukemia (AML). Clinical symptoms in GATA2 patients are diverse and mechanisms driving GATA2 related phenotypes largely unknown. To explore the impact of GATA2 haploinsufficiency on hematopoiesis, we generated a zebrafish model carrying a heterozygous mutation of gata2b (gata2b^+/−^), an orthologue of GATA2. Morphological analysis revealed myeloid and erythroid dysplasia in gata2b^+/−^ kidney marrow (KM). single nucleus (sn)-ATAC-seq showed that the co-accessibility between the transcription start site (TSS) and a +3.5-4.1kb enhancer was more robust in gata2b^+/−^ zebrafish HSPCs compared to wild type, increasing gata2b expression. This is suggestive of an auto-regulatory feedback mechanism, where gata2b expression remains at sufficient levels after the loss of a single allele to maintain the HSPC pool. As a result, gata2b^+/−^ chromatin is also more accessible in the erythroid and myeloid lineage, causing several defects. scRNA-seq data revealed a differentiation delay in erythroid progenitors, hallmarked by downregulation of intrinsic signals like cytoskeletal transcripts, aberrant proliferative signatures, and downregulation of Gata1a, a master regulator of erythropoiesis, likely preceding erythroid dysplasia. This shows that the cell intrinsic compensatory mechanisms for the maintenance of normal levels of Gata2b to maintain HSPC integrity result in aberrant lineage differentiation and a preleukemia syndrome.

## Introduction

The transcription factor GATA2 plays a major role in the generation and maintenance of the hematopoietic system^1–3^. In humans, heterozygous germline mutations in GATA2 often lead to a loss of function of one allele, causing GATA2 haploinsufficiency. The clinical manifestations of GATA2 haploinsufficiency are broad and include immunodeficiency, pulmonary-, vascular- and/or lymphatic dysfunctions and a strong propensity to develop myelodysplastic syndromes (MDS) or acute myeloid leukemia (AML)^4, 5^. These conditions, now collectively referred to as GATA2 deficiency syndromes, were previously known as Emberger syndrome^6^, Monocytopenia and Mycobacterium Avium Complex (MonoMAC) syndrome^7^, dendritic cell, monocyte, B- and natural killer (NK) cell deficiency (DCML)^8^, and familial forms of AML^9^. In addition to the various disease phenotypes, the risk of developing MDS/AML in GATA2 deficient patients is approximately 80% before the age of 40^4, 5^. No clear correlation is found between the occurrence of GATA2 mutations and the severity of hematopoietic deficiencies, even among family members who share the same mutation^10, 11^. Therefore, it is essential to gain insight into the mechanism of underlying GATA2 deficiencies in well-defined uniform experimental models.

In mice, Gata2 has an essential regulatory function in hematopoietic stem cell (HSC) generation and maintenance^1–3^. However, whereas Gata2-null mice are lethal at embryonic day (E) 10.5^3^, Gata2 heterozygous (Gata2^+/−^) mice survive to adulthood with normal blood values. Notwithstanding this apparently normal blood phenotype, Gata2^+/−^ mice have a diminished HSC compartment in the bone marrow (BM) showing a reduced repopulation capacity in competitive transplantation studies^12, 13^. Whereas mouse models thus emerged as a useful source to identify the function of GATA2 in HSC generation and fitness, they leave the mechanisms causing the different aspects of GATA2 deficiency syndromes largely undiscovered.

To better understand the biology of GATA2 haploinsufficiency syndromes, zebrafish serves as an attractive alternative model. Zebrafish have the advantage of having two GATA2 orthologues; Gata2a and Gata2b. Gata2a is expressed predominantly in the vasculature^14^ and is required for programming of the hemogenic endothelium^15, 16^. Gata2b is expressed in hematopoietic stem/progenitor cells (HSPCs)^14^ and homozygous deletion (gata2b^-/−^) redirects HSPC differentiation to the lymphoid lineage in expense of myeloid differentiation causing a lymphoid bias with an incomplete B-cell differentiation in the kidney marrow (KM), thus mimicking one of the GATA2 haploinsufficiency phenotypes found in patients. Additionally, the most primitive HSC compartment was lost in gata2b^-/−^ KM, but none of the animals displayed signs of dysplasia^16, 17^. Because patients carry heterozygous rather than homozygous GATA2 mutations, we specifically focused on how Gata2b haploinsufficiency could be mechanistically linked to erythro-myelodysplasia, the major clinical hallmark of GATA2 patients.

To validate the model, we first assessed hematopoietic cell differentiation in gata2b heterozygous zebrafish (gata2b^+/−^) KM and observed erythroid dysplasia in all gata2b^+/−^ but not in wild-type (WT) zebrafish and myeloid dysplasia in 25% of the gata2b^+/−^ zebrafish. Subsequent single-cell RNA and ATAC sequencing analysis revealed an auto-regulatory feedback mechanism of Gata2b where gata2b chromatin was over-accessible, underlying the hematopoietic lineage differentiation defects in adults. Further characterization of gata2b^+/−^ zebrafish showed reduction of HSC proliferation, failure to initiate the “GATA2 to GATA1 switch” and depletion of Gata1a and its co-factor FOG1 in erythroid progenitor cells. Taken together, these results reveal a dosage-dependent function of Gata2b and provide a plausible explanation for the hematopoietic defects observed in GATA2-deficient patients.

## Material and methods

### Generation and genotyping of Gata2b heterozygous zebrafish

Gata2b^+/−^ and wild type (WT) clutch mates were used for all analyses^16^ and animals were maintained under standard conditions. A knockout allele was generated by introducing a 28bp out-of-frame insertion in exon 3 as previously described^16^.

Zebrafish embryos were kept at 28.5°C on a 14h/10h light-dark cycle in HEPES-buffered E3 medium. Zebrafish were anesthetized using tricaine and euthanized by ice-water. Animal studies were approved by the animal Welfare/Ethics Committee in accordance to Dutch legislation.

### Kidney marrow isolation and analysis

Kidney marrow was dispersed mechanically using tweezers and dissociated by pipetting in phosphate buffered saline (PBS)/10% fetal calf serum (FCS) to obtain single-cell suspensions as previously described^16^. The KM cells were sorted in non-stick cooled micro tubes (Ambion) containing 10% FCS in PBS. Proliferation was assessed by anti-Ki67 staining in fixed (4% paraformaldehyde) KM cells. 7-AAD (7-amino-actinomycin D) (Stem-Kit Reagents) 0.5mg/L or DAPI 1mg/L was used for live/dead discrimination. FACS sorting and analysis were performed using FACS AriaIII (BD Biosciences).

### May–Grünwald-Giemsa stain of KM smears

Kidney marrow smears were fixed in 100% MeOH before staining in May-Grünwald solution (diluted 1:1 in phosphate buffer) and Giemsa solution (diluted 1:20 in phosphate buffer) followed by a last rinsing step in tap water.

Morphological analysis was performed by pathologist by counting 200-500 hematopoietic cells of each kidney marrow smear; excluding mature erythrocytes and thrombocytes. Cells were categorized as: blast, myelocyte, neutrophil, eosinophil, lymphocyte or erythroblast. Furthermore, if dysplasia was observed within a specific lineage, the percentage of dysplastic cells within that lineage was determined by additional counting of at least 50 cells within that specific lineage.

### Single cell omics sequencing

For single cell transcriptomics sequencing (scRNA-seq), kidney marrow cells were isolated and 7×10^4^ viable cells were sorted from 2 pooled Tg(CD41:GFP^18^; runx1:DsRed^19^) WT or gata2b^+/−^ male zebrafish at 1 year of age. For additional replicates, 7×10^4^ single viable cells were sorted from kidney marrows of one WT and one gata2b^+/−^ female zebrafish between 18-20 months of age. cDNA was prepared using the manufacturers protocol (Chromium Single Cell 3’ reagent kits kit v3, 10x Genomics) and sequenced on a Novaseq 6000 instrument (Illumina). After sample processing and quality control analysis, 18,147 cells for WT and 10,849 cells for gata2b^+/−^ were processed for further analysis. The read depth was over 20K reads for all replicates.

We collected the same KM cell composition as scRNA-seq for single nucleus chromatin accessibility sequencing (snATAC-seq, Chromium Single Cell ATAC reagent kits, 10x Genomics). After low quality data filtering, 4,190 nuclei for WT and 7,572 nuclei for gata2b^+/−^ were processed for downstream analysis.

Data was analyzed using the Seurat^20^ and Signac^21^ pipeline. Trajectory inference was performed using Monocle3^22^ R packages.

### Statistics

Statistical analysis was carried out in GraphPad Prism 8 (GraphPad Software). Unless otherwise specified, data were analyzed using unpaired, 2-tailed Student’s t-test. Statistical significance was defined as p<0.05. Graphs are means ± standard error of mean (SEM) and the number of replicates is indicated in the figure legend.

## Results

### Gata2b^+/−^ zebrafish have erythroid and myeloid dysplasia in the kidney marrow

We first assessed hematopoietic cell morphology in KM smears of WT and gata2b^+/−^ zebrafish with ages ranging from 9 months post fertilization (mpf) to 18 mpf. Morphological analysis showed that, while WT zebrafish had normal KM cell morphology, all gata2b^+/−^ KM samples had a considerable fraction of dysplastic cells in the erythroid lineage (Figure 1A, panel 1-6). On average 0.5% of WT erythroid cells showed dysplastic features, compared to 9.9% of gata2b^+/−^ erythroid cells (Figure 1A and B), the latter representing 4.5% of the total kidney marrow population of gata2b^+/−^ zebrafish. Myeloid lineage dysplasia was seen in the gata2b^+/−^ KM in 25% of the fish (Figure 1A, panel 7 and 8, and B). In these samples, 30% of myeloid cells were dysplastic compared to 0.3% in WT. Myeloid dysplasia was mostly represented by multi-lobulated nuclei, whereas the erythroid abnormalities ranged from nuclear deformities and double nuclei to irregular cytoplasm or an almost complete lack of cytoplasm (Figure 1A). The remaining cell types were not affected morphologically by Gata2b haploinsufficiency. These results indicate that gata2b heterozygosity induces dysplasia, predominantly in erythroid and myeloid progenitors.

**Figure. 1:**
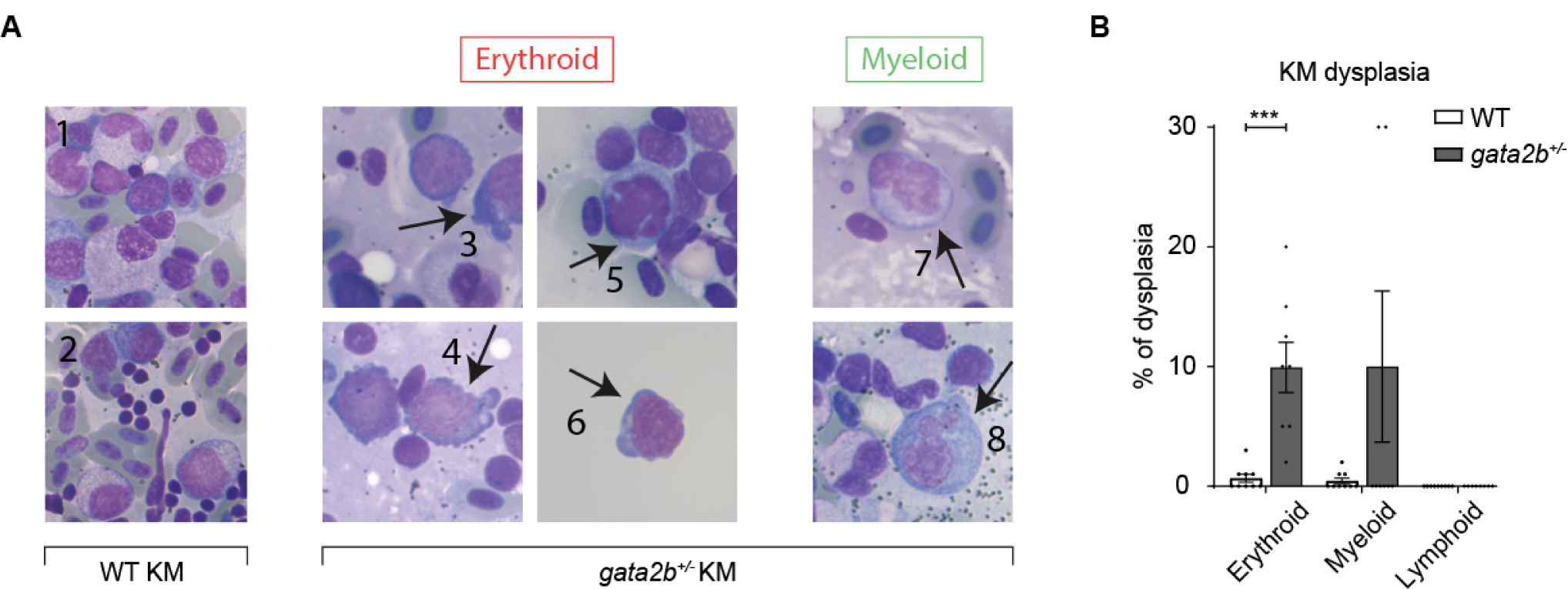
Gata2b^+/−^ kidney marrow shows erythroid and myeloid dysplasia (A) Representative pictures of kidney marrow smears after May–Grünwald-Giemsa staining of WT KM smears (panel 1 and 2) and gata2b^+/−^ KM smears (panel 3-8). 3) Blebbing in cytoplasm of proerythroblast; 4) Irregular cytoplasm in erythroid precursor; 5) Lobed nucleus and micronucleus in erythroblast; 6) Blebbing in cytoplasm of blast of sorted cell after cytospin; 7) Binucleated promyelocyte; 8) Multinucleated promyelocyte. (B) Frequency of dysplastic cells of the erythroid, myeloid and lymphoid lineage in KM smears of WT (n=8) and gata2b^+/−^ (n=8) zebrafish. ***: P < 0.001

To investigate if hematopoietic lineage differentiation ratios were still intact, WT and gata2b^+/−^ KM was assessed by flow cytometry (supplemental Figure 1A). Gata2b^+/−^ zebrafish KM showed no significant difference in the distribution of either mature myeloid-, erythroid-, lymphoid- or progenitor- and HSPC populations compared to WT (supplemental Figure 1B-E). Since GATA2 haploinsufficiency manifestations might require longer periods of time to become evident^4, 5^, we tested the effect of gata2b heterozygosity in zebrafish up to 18 months of age. However, no significant differences were found in the ratios of different lineages in gata2b^+/−^ kidney marrow compared to WT during this period (supplemental Figure 1B-E). Interestingly, the erythroid population significantly increased, indiscriminately of genotype, after 12 months of age (P<0.001) (supplemental Figure 1D), while the remaining myeloid, lymphoid and HSPC and progenitor populations did not vary as dramatically (supplemental Figure 1B, C and E), suggesting aging results in an erythroid biased hematopoietic system in zebrafish. To confirm the presence of a normal hematopoietic lineage distribution in gata2b^+/−^ kidney marrow we bred Gata2b-zebrafish with transgenic GFP reporter zebrafish. Lineage analysis in transgenic lines specifically marking neutrophils by Tg(mpx:GFP) (supplemental Figure 2A-B), T-cells by Tg(lck:GFP) (supplemental Figure 2C-D), B-cells by Tg(Igm:GFP) (supplemental Figure 2E-F) and Tg(mpeg:GFP) (supplemental Figure 2G-H) or monocytes by Tg(mpeg:GFP)^23, 24^ (supplemental Figure 2I-J) showed no significant alterations in lineage distribution between WT and gata2b^+/−^ KM ^25–28^. KM smear quantification by May–Grünwald-Giemsa (MGG) staining revealed that gata2b^+/−^ KM had a significant decrease in eosinophils compared to WT (supplemental Figure 2K), representing less than 5% of the total KM cells. In summary, based on flow cytometry analysis and transgenic reporters we conclude that the differentiation in major hematopoietic lineages is not altered in gata2b^+/−^ KM and that the erythroid and myeloid dysplasia does not affect this.

### Single cell RNA- and ATAC sequencing analysis reveals a cell maturation delay in gata2b^+/−^ KM

To investigate the molecular mechanisms underlying dysplasia in gata2b^+/−^ KM, we assessed chromatin accessibility, a measure for gene activation, and transcriptional differences in dysplastic cell populations. We flow-sorted 4 cell populations based on light scatter and observed dysplastic cells in the progenitor and lymphoid + HSPCs population of gata2b^+/−^ KM, indicating that dysplastic cells could be viably sorted (Figure 1A, panel 6). We did not identify a uniquely separated population of dysplastic cells, possibly caused by a heterogeneity in their shape. Therefore, we sorted progenitor and lymphoid + HSPC populations from WT and gata2b^+/−^ KMs and performed single-cell (sc) RNA- and single nucleus (sn) ATAC-sequencing to identify transcriptome and epigenetic regulation signatures defining dysplastic cells (Figure 2A). After filtering out low quality cells, we obtained 28,996 cells (WT = 18,147, gata2b^+/−^ = 10,849) for scRNA-seq analysis and 11,762 cells (WT = 4,190, gata2b^+/−^ = 7,572) for snATAC-seq analysis (Figure 2B).

**Figure. 2:**
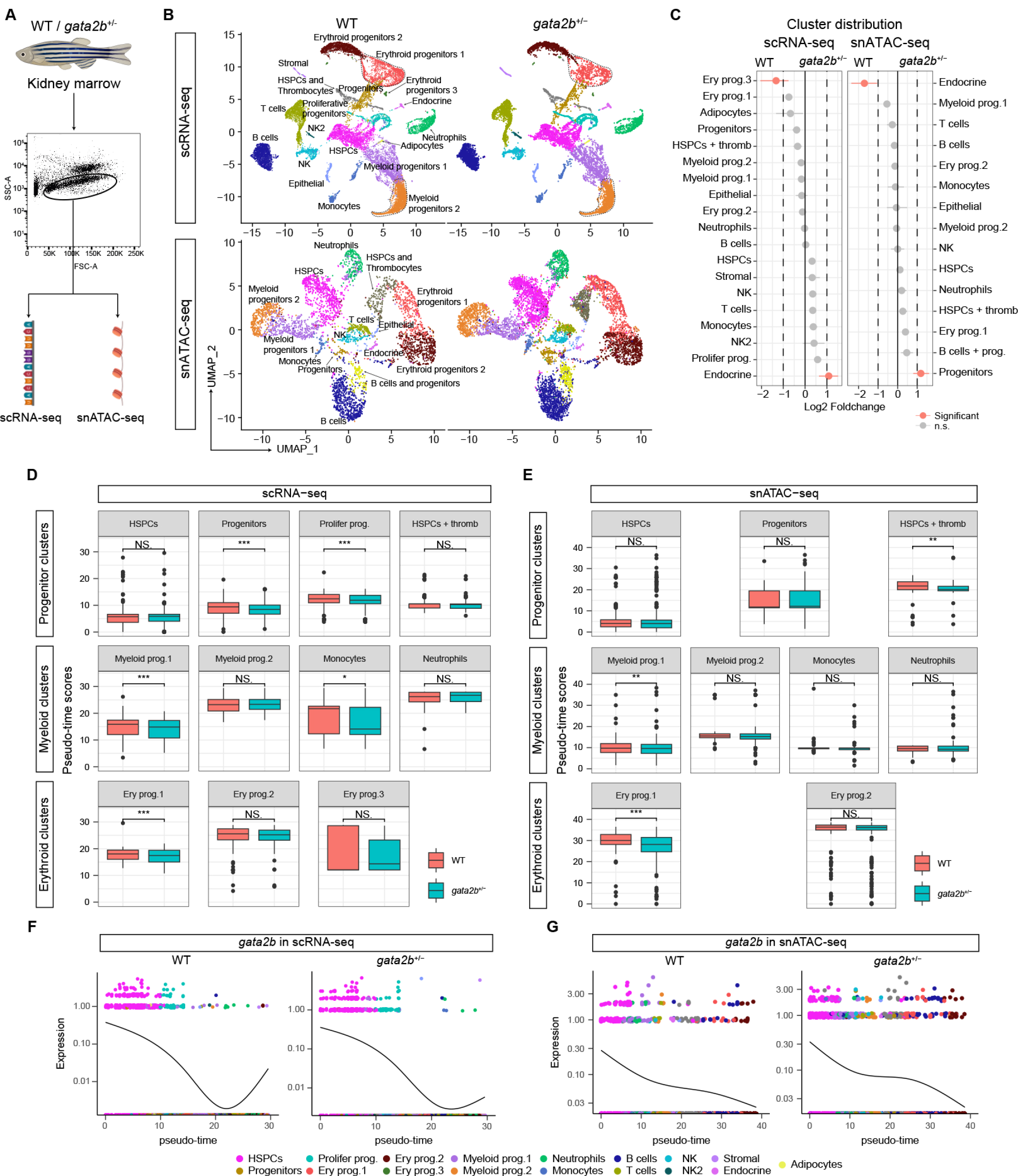
Single cell transcriptome and chromatin accessibility analysis reveals a differentiation block in gata2b^+/−^ KM (A) The flow cytometry-based sorting strategy for kidney marrow cells for scRNA-seq and snATAC-seq analysis. (B) UMAP analysis of scRNA-seq (top panels) and snATAC-seq (bottom panels) showing all cell types, which was 19 cell types in scRNA-seq and 16 cell types in snATAC-seq. UMAP atlas was separated between WT and gata2b^+/−^ cells. (C) Cluster distribution quantification of scRNA-seq (left panel) and snATAC-seq (right panel) between WT and gata2b^+/−^ cells, with significantly differentially distributed clusters represented in pink. A statistical significance threshold of FDR < 0.05 and log2 fold change > 1 were applied to determine the significance of the observed differences. (D-E) Box plots representing the comparison of pseudo-time scores across progenitor, myeloid, and erythroid clusters in D) scRNA-seq data and E) snATAC-seq data. *: P < 0.05, **: P < 0.01, ***: P < 0.0001, NS: not significant. (F) WT (left panel) and ^+/−^ (right panel) expression of gata2b in scRNA-seq data in pseudo-time. Dots are depicted if gata2b expression is detected and colored according to cell type. The curved line shows the dynamic expression of gata2b throughout pseudo-time trajectory. (G) WT (left panel) and gata2b^+/−^ (right panel) chromatin accessibility of the gata2b locus in snATAC-seq data in pseudo-time. Dots are depicted if open chromatin was detected and are colored according to the cell types. The curved line shows the open chromatin at the gata2b locus throughout pseudo-time trajectory.

scRNA-sequencing resulted in 19 clusters based on a nearest neighbor algorithm using the R Seurat package^20^ (Figure 2B). Each cluster was classified based on differentially expressed genes (supplemental Figure 3A-D) and known differentiation markers^29–34^ (Figure 2C). We then used this labeled scRNA-seq dataset as a reference to interpret and identify 16 clusters in the snATAC-seq dataset based on the R Signac package^21^ (Figure 2B, supplemental Figure 4A-C). Cluster proportion analysis indicated an overall similar distribution of cells within clusters between gata2b^+/−^ and WT (Figure 2C). However, a clear difference in differentiation trajectory was observed between WT and gata2b^+/−^ KM in scRNA-seq and snATAC-seq data based on the R package Monocle3^22^, and particularly note the difference in pseudo-time scores between WT and gata2b^+/−^ erythroid progenitors 1 and myeloid progenitors 1, suggesting a delayed differentiation in gata2b^+/−^ KM (Figure 2D and E, supplemental figure 4D). Whereas the pseudo-time score for HSPCs was similar between WT and gata2b^+/−^ zebrafish, the pseudo-time scores of differentiation of erythroid and myeloid lineages were reduced in gata2b^+/−^ KM (Figure 2D and E). Moreover, the expression of gata2b was reduced in gata2b^+/−^ cells compared to WT in the terminal trajectory of hematopoietic development, where erythroid and myeloid progenitors are located (Figure 2F and G). Based on these findings, we conclude that gata2b^+/−^ KM showed an overall lineage maturation delay. As expected, a clear distinction in open chromatin patterns of lineage specific transcription factors (TF) between immature and mature cells was seen in WT KM (Supplemental figure 5A, C and E, left panels). In contrast, this distinction was less pronounced in gata2b^+/−^ KM (Supplemental figure 5B, D and F, left panels). Specifically, the ETV family of transcription factors and the GATA family transcription factors were affected in erythroid differentiation, SPI1, SPIB, and CEBPA family of transcription factors were affected in myeloid differentiation, and TCF4 was most affected in lymphoid differentiation and showed significantly different patterns of activation (Supplemental figure 5A-F). Surprisingly, this did not always lead to a reduction in gene expression (Supplemental figure5A-F, right panels), particularly note the increased expression of gata2b in erythroid lineage differentiation (Supplemental figure 5B, right panel).

### gata2b heterozygosity results in over-accessible gata2b chromatin leading to self-renewal defects in HSPCs

To further investigate the mechanism underlying the effects of haploinsufficient gata2b expression, we assessed the accessible peak distribution of gata2b and its upstream region. First of all, the chromatin accessibility signals of gata2b were strongest in the HSPCs cluster (Figure 3A, top line), then gradually disappeared with cell maturation (Figure 3A, all other lines), which is consistent with the GATA2 function in HSC maintenance^1–3^. However, when we compared the signals between WT on the left and gata2b^+/−^ on the right, we observed that the chromatin of the gata2b gene body and upstream regions was more accessible in gata2b^+/−^ HSPCs. Chromatin co-accessibility analysis, which was used to predict cis-regulatory interaction in the genome, suggested that there were two regions, +3.5-4.1kb termed enhancer 1(E1) and +5.4-6.2kb, termed enhancer 2 (E2), which could be strongly interacting with each other (Figure 3A, black arrows in Links). Moreover, E1 had stronger interaction with gata2b transcription start site (TSS) in gata2b^+/−^ KM (Figure 3A, red arrows in Links). These two upstream regions might work as enhancers for gata2b transcription. When we assessed the cell distribution of the two enhancers and gata2b TSS (Figure 3B-G), we observed that the accessibility signals of +5.4-6.2kb region-E2, (Figure 3B, C), was accessible in HSPCs and gata2b^+/−^ erythroid progenitors, while a +3.5-4.1kb region-E1, (Figure 3D, E) was mainly accessible in HSPCs and gata2b^+/−^ myeloid progenitors. This result suggests that E1 and E2 might be responsible for different functions in hematopoiesis, and particularly in gata2b^+/−^ KM, E1 affects myeloid differentiation and E2 affects erythroid differentiation. The percentage of cells with gata2b mRNA expression did not show a significant difference as a result from the up-regulated chromatin accessibility and stronger enhancer-TSS co-accessibility (Supplementary figure 6A and B), but we did see an increase in the level of gata2b expression in the HSPC cluster as a result of the increase in enhancer-TSS co-accessibility (Figure 3A, H, and I and Supplementary Figure 5B, right panel).

**Figure. 3:**
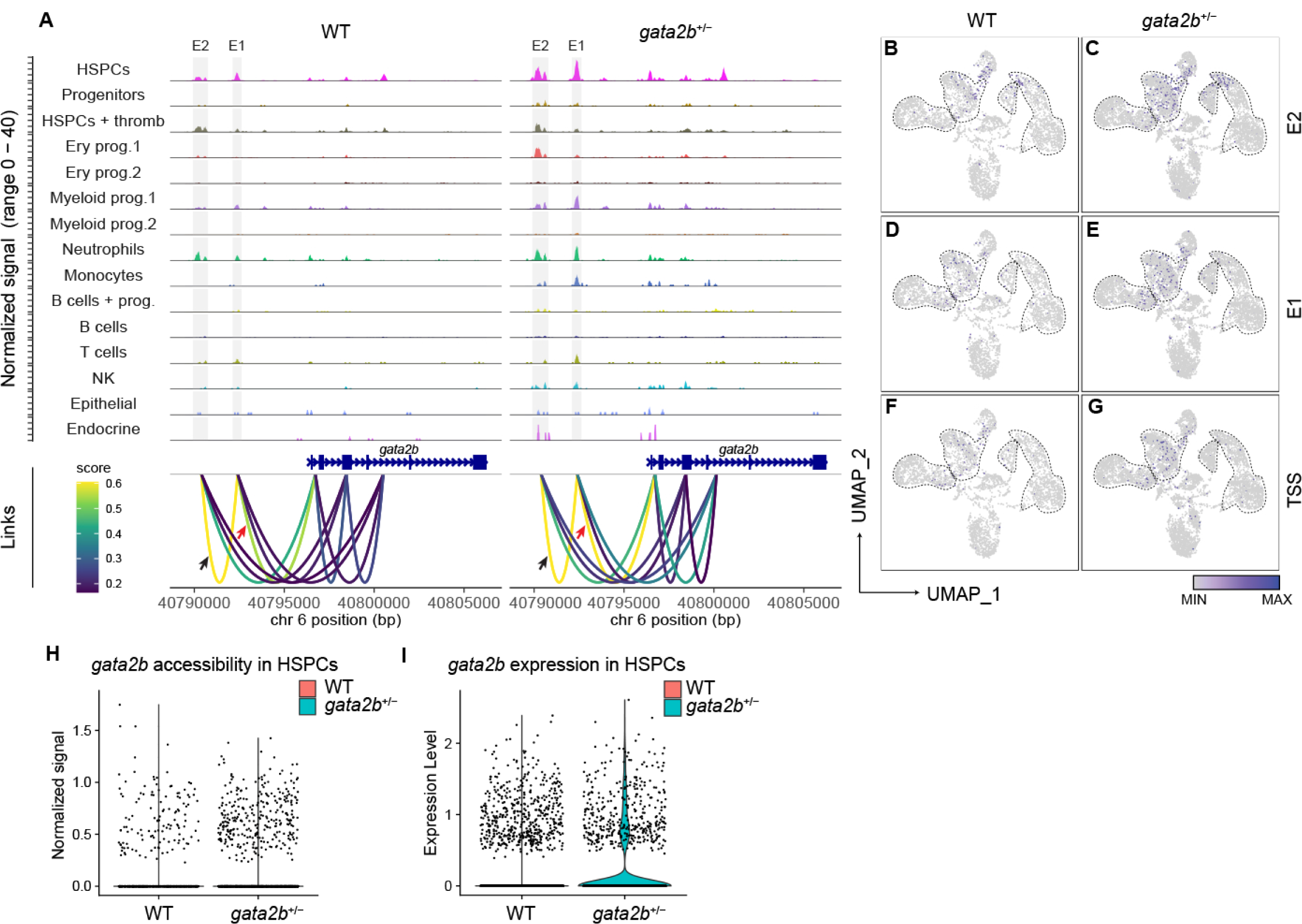
Differences in chromatin accessibility of Gata2b result in upregulation of Gata2b expression in gata2b^+/−^ HSPCs (A) Coverage plots showing significant differences of gata2b chromatin accessibility between WT and gata2b^+/−^ cells with the location of the gata2b gene block in blue. The normalized peak signal range is 0 to 40. Highlighted genomic regions within grey blocks indicate the location of two enhancers, enhancer 1 (E1) and enhancer 2 (E2). The link between the blocks illustrate predicted cis-regulatory interactions between the gata2b gene body and its upstream region, based on examination of genome co-accessibility. Link colors correspond to predicted co-accessibility scores, with yellow indicating stronger predicted interactions. Black arrows indicated the co-accessibility between E2 and E1. Red arrows indicated the co-accessibility between E1 and TSS which is stronger in gata2b^+/−^ KM. (B-G) UMAP feature plots depict the open chromatin of Enhancer 2 (E2; B and C), Enhancer 1 (E1; D and E), and transcription start site (TSS; F and G) of gata2b. UMAP atlas were separated by WT (B, D and F) and gata2b^+/−^ cells (C, E and G). (H) Open chromatin of the gata2b locus per HSPCs cluster cell comparing WT and gata2b^+/−^HSPCs. (I) Violin plot displaying the expression level of gata2b in HSPCs in WT (pink) and gata2b^+/−^ (turquois).

### The functional consequence of epigenetic changes of the gata2b locus in HSPCs is a reduction in G2-M phase in cell cycle

Because the most significant signal differences occurred in HSPC clusters and the cell differentiation velocity differences appeared at “progenitors” clusters (Figure 2D), we compared the different TF activity in progenitor-like clusters, including “HSPCs”, “progenitors”, and “HSPCs and thrombocytes”. Gata2b^+/−^ progenitor-like clusters showed significantly up-regulated chromatin accessibility of motifs of the BATF family of transcription factors (Figure 4A). The BATF family of TFs are known to function in limiting the self-renewal of HSCs in response to the accumulation of DNA damage^35^, suggesting a defect in self-renewal of gata2b^+/−^ HSCs. To investigate this, proliferation was assessed in CD41:GFP^int^ purified HSCs (Figure 4B). Although numerically unaffected by gata2b heterozygosity (Figure 4C), cell cycle phase distribution was markedly different in gata2b^+/−^ HSCs compared to WT (Figure 4D and E), with increased G1 phase and decreased G2-M phase. As stem cells are known to have a short G1 phase to reduce cellular differentiation potential, this may already be an indication that these HSCs lose self-renewal capacity and differentiate, which is halted in the next differentiation stages^36^.

**Figure 4.**
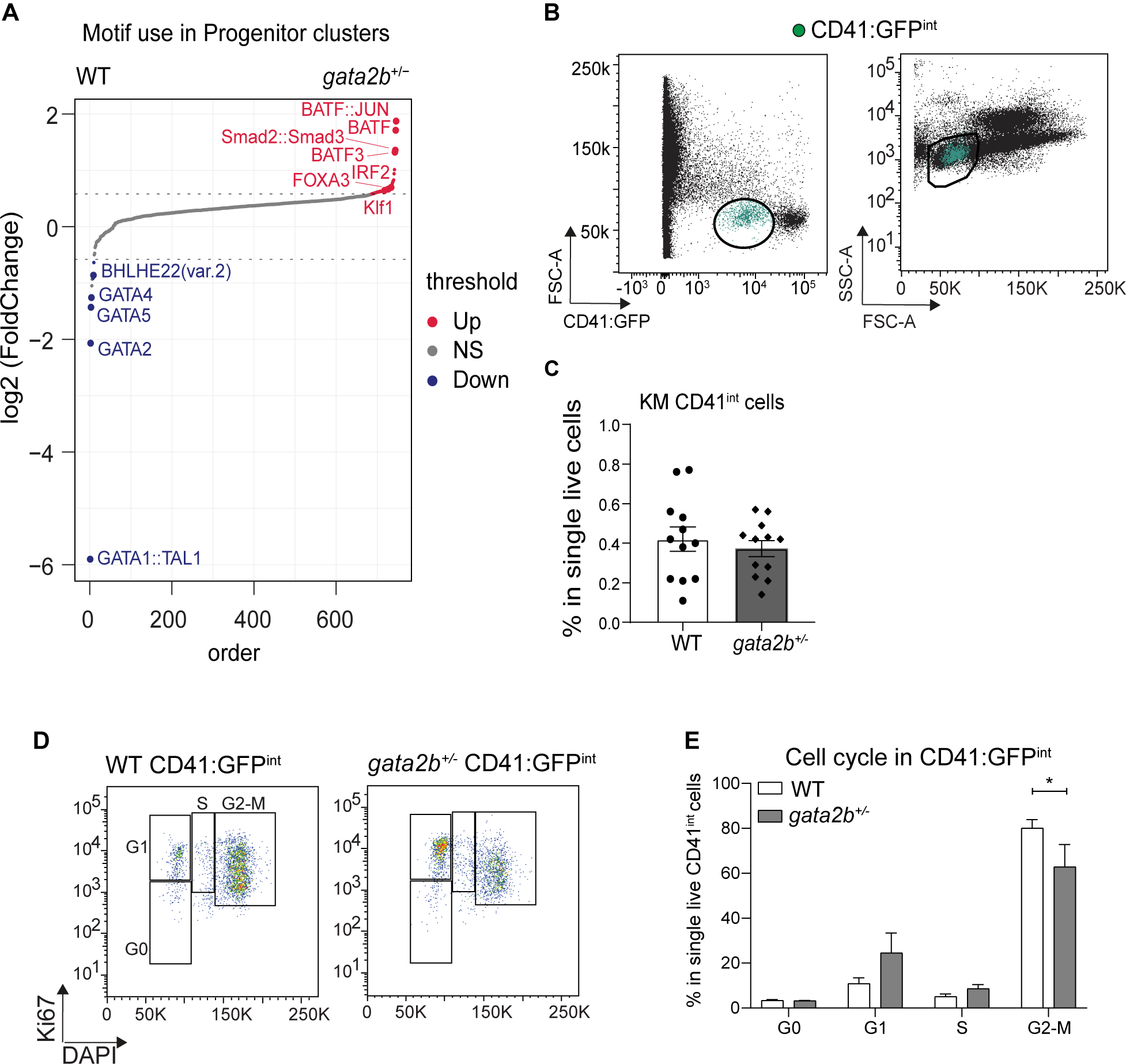
Functional consequences in HSPCs upon Gata2b overexpression. (A) The dot plot displays the log2 fold change values of differentially activated transcription factor (TF) motifs in progenitor clusters, including "HSPCs", "progenitors", and "HSPCs and thrombocytes", between WT and gata2b^+/−^ cells. The TF motifs are ranked in increasing order based on log2 fold change values. Dotted lines indicate the threshold for significance, with absolute log2 fold change values > 0.58. The blue and red colors represent TF motifs that are statistically significant (P value < 0.05) with absolute log2 fold change values > 0.58. (B) Representative figure of identification of CD41:GFP^int^ population (in green) and distribution in the FSC-A SSC-A kidney marrow population. (C) Quantification of the percentage of CD41:GFP^int^ cells in WT (n=12) and gata2b^+/−^ (n=12) KM single live cells. (D) Representative flow cytometry plots of cell cycle analysis by anti-Ki67 and DAPI staining in WT and gata2b^+/−^ CD41:GFP^int^ KM. (E) Quantification of the percentages of WT (n=3) and gata2b^+/−^ (n=3) CD41:GFP^int^ cells in different cell cycle stages (* P value<0.05). Data represents as mean ± standard error of the mean.

Epigenetic dysregulation results in delayed erythroid maturation in gata2b^+/−^ kidney marrow Next, we sought to reveal the molecular mechanism behind the erythroid dysplasia in Gata2b^+/−^ KM. Compared to WT, Gata2b^+/−^ progenitor-like clusters had significantly reduced accessibility of GATA2 and GATA1::TAL1 motifs (Figure 4A). Despite the increase in gata2b expression, this indicated an overall reduction of Gata2b function in activation of downstream targets in the gata2b^+/−^ zebrafish genome. Activation of GATA2 motifs were mainly enriched in the erythroid progenitors (Figure 5A). An important downstream target of Gata2 is Gata1, which is the master regulator of erythropoiesis^37, 38^. We therefore investigated Gata1a accessibility and expression in our dataset (Figure 5B-E). The gata1a TSS was generally more accessible in gata2b^+/−^ KM (Figure 5B), but not in erythroid progenitors 2 (Figure 5C). Furthermore, we found downregulation of gata1a mRNA expression (Figure 5D) and the GATA1::TAL1 motif was less accessible (Figure 5E), further supporting the reduced functionality of Gata1a in erythroid progenitors. This could be due to reduced expression of zfpm1 (also known as Friend of GATA (FOG1)), which can facilitate GATA1 expression^39^(Figure 5F). The increased expression of gata2b in HSPCs (Figure 3I) and reduced expression of gata1a in erythroid progenitors (Figure 5D) suggested that the “GATA factor switch”, which is a shift from GATA2 to GATA1 occupation during erythroid lineage differentiation^40–42^, might be silenced, causing deficiencies in erythroid lineage differentiation.

**Figure. 5:**
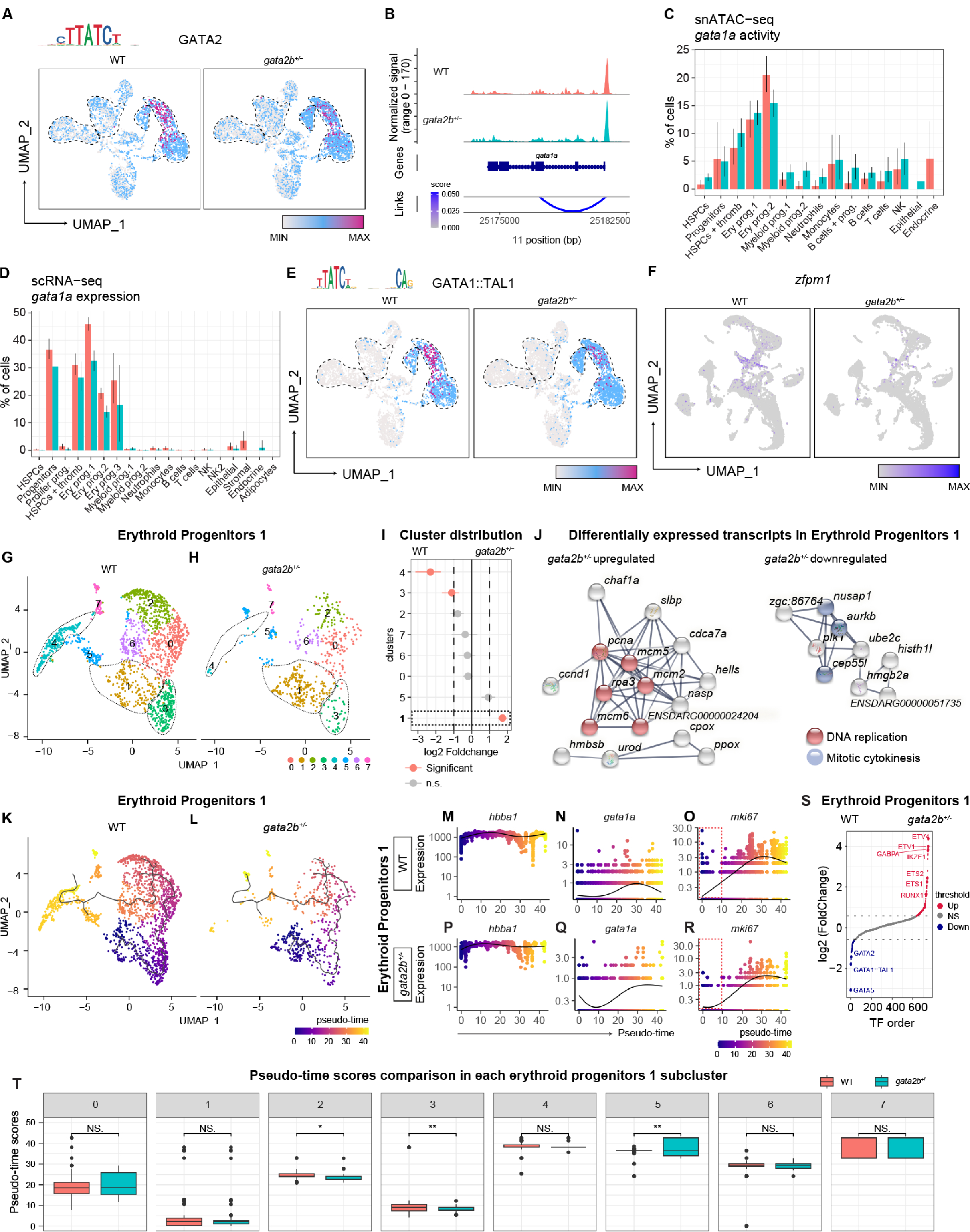
Epigenetic changes resulting in the erythroid differentiation block in gata2^+/−^ KM (A) UMAP feature plots depicting open chromatin of GATA2 motif sequence indicating activity of GATA2. UMAP atlas were separated by WT (left panel) and gata2b^+/−^ cells (right panel). The top left corner of UMAP plots display the TF sequence logo corresponding to the analyzed motif, generated by the function MotifPlot. (B) Coverage plots showing differences of gata1a chromatin accessibility between WT and gata2b^+/−^ cells. (C) Percentage of cells derived from WT and gata2b^+/−^ KM with accessibility of the gata1a locus across clusters in snATAC-seq data. (D) Percentage of cells derived from WT and gata2b^+/−^ KM expressing gata1a across clusters in scRNA-seq data. (E) UMAP plots depict the activity of combined motif GATA1::TAL1. UMAP atlas were separated by WT (left panel) and gata2b^+/−^ cells (right panel). The top left corner of the UMAP plots display the TF sequence logo corresponding to the analyzed motif, generated by the function MotifPlot. (F) Feature plot showing the zfpm1 (FOG1) expression in WT and gata2b^+/−^ KM. (G-H) UMAP plots of the subclustering of the erythroid progenitors 1 cluster in scRNA-seq data. UMAP atlas were separated by WT(G) and gata2b^+/−^ cells (H) (I) Quantification of distribution between WT and gata2b^+/−^ erythroid progenitor 1 subclusters. Significantly differentially distributed clusters in pink. Dotted box indicates overrepresented subcluster in gata2b^+/−^ cells. (J) STRING network of upregulated and downregulated transcripts in gata2b^+/−^ “erythroid progenitors 1”. Only networks with more than 2 interactions were represented. Highlighted in red DNA replication genes from KEGG pathways and highlighted in blue mitotic cytokinesis genes from biological processes (Gene Ontology). (K-L) Lineage trajectory analysis of WT (K) and gata2b^+/−^(L) erythroid progenitor 1 subclusters. (M-R) WT (M-O) and gata2b^+/−^ (P-R) pseudo-time expression of individual genes hemoglobin beta adult 1 (hbba1) (M and P), GATA binding protein 1 alpha (gata1a) (N and Q), and mki67 (O and R). Dotted red boxes indicate the different initial expression of mki67 (O and R). Fold change > 0.05 & adjusted P value < 0.05. FDR=False discovery rate. FD=Fold difference. (S) Dot plot displaying log2 fold change values of differentially activated TF motifs in the erythroid progenitor 1 cluster between WT and gata2b^+/−^ cells in snATAC-seq data. The TF motifs are ranked in descending order based on log2 fold change values. Dotted lines indicate the threshold (absolute log2 fold change < −0.58 and >0.58) for significance, The blue (downregulated) and red (upregulated) colors represent TFs that are statistically significant (P value < 0.05). (T) Box plots representing the comparison of pseudo-time scores between WT and gata2b^+/−^across all subclusters in erythroid progenitors 1. *: P < 0.05, **: P < 0.01, NS: no significance.

Considering that the dysplastic cells in KM smears were of myeloid or erythroid origin, we hypothesized that dysplastic cells occupy a proportion of erythroid or myeloid clusters. For a more precise examination of small populations along the myeloid and erythroid differentiation lineage, we sub-clustered ‘’HSPCs’’, ‘’proliferative progenitors’’, “progenitors”, “myeloid progenitors 1”, “myeloid progenitors 2”, “erythroid progenitors 1”, and “erythroid progenitors 2” clusters and identified 5 significantly overrepresented sub-clusters in gata2b^+/−^ KM (Figure 5G-I, supplemental Figure 7 A-H). These sub-clusters represented 5.6% of the total sequenced gata2b^+/−^cells, comparable to the proportion of dysplastic cells observed in gata2b^+/−^ KM. Differentially expressed genes (DEGs) analysis of overrepresented sub-clusters in gata2b^+/−^ KM compared to the rest of that cluster showed downregulation of tubulin transcripts (not shown), suggesting a loss of cytoskeletal structure, a characteristic of dysplasia. Besides, TF motif score analysis with snATAC-seq indicated a general reduction for lineage-specific determination and differentiation motifs in gata2b^+/−^ KM (supplemental Figure 5A-F), suggesting that gata2b^+/−^ cell maturation could be impaired resulting in a differentiation block, as this is often the case in dysplasia^43^.

To identify the mechanisms behind the erythroid differentiation delay in gata2b^+/−^ KM, we compared the DEGs and their function in the “erythroid progenitors 1” cluster. The result showed that gata2b^+/−^ “erythroid progenitors 1” cluster cells had upregulated signatures of DNA replication together with the downregulation of transcripts necessary for mitosis (Figure 5J), which can explain the origin of multi-lobulated nuclei and other nuclear abnormalities observed in gata2b^+/−^ dysplastic cells^44, 45^. Pseudo-time trajectory inference showed a diminished differentiation in gata2b^+/−^ “erythroid progenitors 1” cluster with a lower pseudo-time score (Figure 5K, L, and T). Pseudo-time dynamic gene expression level also indicated the erythropoiesis differentiation delay in the gata2b^+/−^ “erythroid progenitors 1” cluster with persistent high open chromatin, but lower expression of gata1a, which should gradually decrease toward terminal erythroid maturation, and the decreased initial level of mki67, indicative of incomplete differentiation (comparing Figure 5M-O to Figure 5P-R). This could be due to a diminished open chromatin structure at the locus leading to a reduction in Gata2b binding sites to drive proper Gata1a expression, to support the erythroid differentiation process. Furthermore, TF motif score analysis of the Erythroid progenitors 1 cluster revealed a significant depletion for GATA family motifs, but an enrichment for ETS and ETV family of transcription factor motifs in gata2b^+/−^ “Erythroid progenitors 1” cells (Figure 5S), suggesting an epigenetic dysregulation related to decreased maturation of erythroblasts resulting from Gata2b haploinsufficiency.

## Discussion

In humans, a balanced GATA2 expression is essential for proper hematopoiesis. Consequently, more than 80% of GATA2 mutation carriers progress to hematological malignancy by 40 years of age^4^. While the clinical consequences of GATA2 mutations became obvious over the last decades, the regulation of GATA2 activity and its contribution to human bone marrow failure syndromes and progression to hematological malignancy remain incompletely understood.

Here, we used transgenic reporters, morphological phenotyping, and single-cell omics sequencing to analyze gata2b heterozygous zebrafish. We showed that, while major differentiation lineages remain intact, gata2b heterozygosity causes dysplasia in erythroid and myeloid progenitors of zebrafish KM. snATAC-seq revealed an auto-regulatory feedback mechanism for Gata2b haploinsufficiency, characterized by gata2b chromatin over-accessibility and stronger co-accessibility of gata2b enhancers and TSS, indicating epigenetic dysregulation in hematopoiesis in the zebrafish model. scRNA-seq analysis did not identify a single population of dysplastic cells. Interestingly, a single erythroid progenitor cluster was identified using snATAC-seq, and could indicate a specific dysplastic erythroid population. The difference in recognition of the dysplastic cluster between scRNA-seq and snATAC-seq may point to the underlying mechanism of the dysplasia where we have evidence that the epigenetic compensatory mechanism of maintaining normal levels of Gata2b may cause the dysplastic features during lineage differentiation, particularly in the GATA2 to GATA1 switch and therefore affects erythroid lineage differentiation most. This also underscores the importance of epigenetic analysis in other types of MDS and the fact that many epigenetic regulators are mutated in MDS and AML^46^. In gata2b^+/−^ erythroid progenitors we found increased proliferative signatures together with decreased expression of genes related to mitosis indicating an impaired cell cycle progression. Furthermore, we found the depletion of GATA family factors indicating a failure of the “GATA factor switch” that is indispensable for erythroid lineage differentiation^40–42^. We propose that these alterations could play a role in the onset of dysplasia found in gata2b^+/−^ zebrafish.

Interestingly, whereas gata2b^-/−^ zebrafish display abrogated myeloid lineage differentiation and a bias toward lymphoid differentiation ^16, 17^, gata2b^+/−^ did not simply result in an intermediate phenotype between WT and gata2b^-/−^. Instead, Gata2b haploinsufficiency uniquely caused dysplasia, not observed in gata2b^-/−^, potentially caused by compensatory mechanisms in haploinsufficiency that are otherwise overcome in gata2b^-/−^ zebrafish^47^. This may also be the reason that missense mutations have a different molecular mechanism, because these alleles are not prone to mRNA degradation and may thus not be compensated by epigenetic deregulation. The differences in the homozygous and heterozygous Gata2b knockout phenotype support a role for gene dosage underlying the GATA2 deficiency phenotype, possibly explaining the phenotypic heterogeneity between patients. Since both erythroid and myeloid dysplasia can be observed in GATA2 patients^4^, we propose that the presence of dysplastic cells in gata2b^+/−^ resembles the clinical phenotypes associated with GATA2 heterozygosity. In the future, the isolation of single dysplastic cells could help us to further explore the effect of Gata2 haploinsufficiency in malignant transformation. Nevertheless, it remains to be established how gata2b^+/−^ HSPCs would respond to secondary insults such as infections or severe bleeding.

In conclusion, while the major lineage differentiation remains intact, gata2b^+/−^ zebrafish possess a stressed proliferative HSPC compartment which leads to the generation of erythroid and myeloid dysplastic cells. Taken together, our model provides insights into the consequences of Gata2b dosage, illustrated the alteration of the microenvironment, and reveal how changes in epigenetics affect the outcome in lineage differentiation after Gata2b haploinsufficiency in zebrafish.

## Acknowledgements

We thank Dr Monteiro (University of Birmingham) for careful reading of the manuscript. We thank the Experimental Animal Facility of Erasmus MC for animal husbandry and the Erasmus Optical Imaging Center for confocal microscopy services. This research is supported by the European Hematology Association (junior and senior non clinical research fellowship) (EdP), the Dutch Cancer Foundation KWF/Alpe d’HuZes (SK10321)(EdP), the Daniel den Hoed Foundation for support of the Cancer Genome Editing Center (IT) and the Josephine Nefkens Foundation for purchase of the Chromium 10x (IT).

## Authorship contributions

EdP and EG conceived the study; EG, CK, HdL, JZ, DB, and EB performed experiments; EG, WZ, CK, RH, KG and EdP analysed results; IPT provided resources and EG, WZ, CK and EdP wrote the manuscript and IPT revised the manuscript.

**Supplementary figure. 1: Lineage gating of total KM in WT and gata2b^+/−^ zebrafish**

(A) Gating strategy of flowcytometry analysis of whole kidney marrow of WT and gata2b^+/−^ zebrafish. Percentages represent the average of all zebrafish analyzed per genotype.

(B-E) Quantitation as percentages of the different cell populations in single viable cells of myeloid (B), lymphoid and HSCs (C), erythrocytes (D) and progenitors (E) in WT and gata2b^+/−^ zebrafish kidney marrow over time. Statistical analysis showing as mean ± standard error of the mean. ***: P < 0.001.

**Supplementary figure. 2: Transgenic zebrafish reporter lines show no overall differentiation difference in gata2b^+/−^**

(A) mpx positive cells, depicted in green, with distribution in FSC-A SSC-A graph.

(B) Quantification of the percentage of mpx:GFP+ cells in WT (n = 8) and gata2b^+/−^ (n = 5) kidney marrow single live cells.

(C) Lck positive cells, depicted in green, with distribution in FSC-A SSC-A graph.

(D) Quantification of the percentage of lck:GFP+ cells in WT (n = 4) and gata2b^+/−^ (n = 7) kidney marrow single live cells.

(E) Igm positive cells, depicted in green, with distribution in FSC-A SSC-A graph.

(F) Quantification of the percentage of IgM:GFP+ cells in WT (n = 3) and gata2b^+/−^ (n = 3) kidney marrow single live cells.

(G-J) mpeg positive cells, depicted in green, mark monocytes (G) and phagocytic B-cells (I). Quantification of the percentage of mpeg:GFP+ cells (H and J) in WT (n = 7) and gata2b^+/−^ (n = 7) kidney marrow single live cells.

(K) Frequency of differentiated cell types in KM cells in smears of WT (n=8) and gata2b^+/−^ (n=8) zebrafish. Differentiated erythrocytes were excluded from quantification. *P value<0.5. Data represents mean ± Standard error of the mean.

**Supplementary Figure. 3: Transcriptomic heatmap and the expression of marker genes**

(A) Unbiased heatmap representing the expression level of the top 5 expressed transcripts per cluster. Transcripts highly expressed in multiple clusters were not repeated in the list (like hemoglobin in erythroid clusters). Transcripts identified only by their chromosome location were not included.

(B-D) FeaturePlot gene expression analysis of CD41 (B) in HSPCs and thrombocytes, mki67

(C) in Proliferative progenitors, and pcna (D) predominantly in undifferentiated clusters.

**Supplementary Figure. 4: Expression and accessibility of marker genes used for cluster annotation and the pseudo-time trajectories inference**

(A) Feature plot showing the scRNA-seq expression of marker gene hemoglobin beta adult 1 (hbba1) in the erythroid lineage, granulin1 (grn1) in the myeloid lineage, cluster of differentiation 37 (CD37) in the B cell lineage, and GATA binding protein 2b (gata2b) in HSPCs to show cluster annotation.

(B) Feature plot showing the snATAC-seq gene activity of hbba1 in the erythroid lineage, grn1 in the myeloid lineage, CD37 in the B lineage and gata2b in HSPCs to show cluster annotation.

(C) Coverage plot illustrating the chromatin accessibility peaks of hbba1 in the erythroid lineage, grn1 in the myeloid lineage, CD37 in the B lineage, and gata2b in hematopoietic stem and progenitor cells (HSPCs) to show cluster annotation.

(D) UMAP plot of scRNA-seq data (upper panels) and snATAC-seq data (bottom panels) colored by pseudo-time scores representing cell developmental trajectories split by WT (left panels) and gata2b^+/−^ (right panels) cells. Lower pseudo-time scores correspond to the root of trajectories, while higher pseudo-time scores represent the terminal differentiation trajectories. The lines plotted on the UMAP atlas illustrate the differentiation paths of cells.

**Supplementary Figure. 5: Motif regulation of lineages differentiation along pseudo-time trajectories show reduced differentiation in gata2b^+/−^ KM compared to WT.**

(A) Heatmap plots showing the dynamic motif activity (left panel) and relative gene expression (right panel) of erythroid cells differentiation along pseudo-time trajectory in WT KM. Left panel depicting clusters in snATAC-seq data, including “HSPCs”, “Progenitors”, “HPSCs and thrombocytes”, “Erythroid progenitors 1”, and “Erythroid progenitors 2”, while right panel depicting clusters in scRNA-seq data, including “HSPCs”, “Progenitors”, “Proliferative progenitors”, “HPSCs and thrombocytes”, “Erythroid progenitors 1”, “Erythroid progenitors 2”, and “Erythroid progenitors 3”.

(B) Heatmap plots showing the dynamic motif activity (left panel) and relative gene expression (right panel) during erythroid cell differentiation along pseudo-time trajectory in gata2b^+/−^ KM. The same clusters as those depicted in Figure S5A were used to generate the heatmaps.

(C) Heatmap plots showing the dynamic motif activity (left panel) and relative gene expression (right panel) during myeloid cells differentiation along pseudo-time trajectory in WT KM. Left panel depicting clusters in snATAC-seq data, including “HSPCs”, “Progenitors”, “HPSCs and thrombocytes”, “Myeloid progenitors 1”, “Myeloid progenitors 2”, “Neutrophils”, and “Monocytes”, while right panel depicting clusters in scRNA-seq data, including “HSPCs”, “Progenitors”, “Proliferative progenitors”, “HPSCs and thrombocytes”, “Myeloid progenitors 1”, “Myeloid progenitors 2”, “Neutrophils”, and “Monocytes”.

(D) Heatmap plots showing the dynamic motif activity (left panel) and relative gene expression (right panel) during myeloid cell differentiation along pseudo-time trajectory in gata2b^+/−^ KM. The same clusters as those depicted in Figure S5C were used to generate the heatmaps.

(E) Heatmap plots showing the dynamic motif activity (left panel) and relative gene expression (right panel) during lymphoid cell differentiation along pseudo-time trajectory in WT KM. Left panel depicting clusters in snATAC-seq data, including “HSPCs”, “Progenitors”, “HPSCs and thrombocytes”, “B cells and progenitors”, “B cells”, “T cells”, and “NK”, while right panel depicting clusters in scRNA-seq data, including “HSPCs”, “Progenitors”, “Proliferative progenitors”, “HPSCs and thrombocytes”, “B cells”, “T cells”, “NK”, and “NK2”.

(F) Heatmap plots showing the dynamic motif activity (left panel) and relative gene expression (right panel) during lymphoid cells differentiation along pseudo-time trajectory in gata2b^+/−^ KM. The same clusters as those depicted in Figure S5E were used to generate the heatmaps.

**Supplementary Figure 6. Cell percentages with snATAC activity of the gata2b locus and gata2b expression per cluster.**

A) Cell percentages with detectable open chromatin of the gata2b locus per cluster. B) Cell percentages with detectable gata2b expression per cluster.

**Supplementary Figure 6. gata2b^+/−^ cells have a different distribution of subclusters compared to WT**

Split UMAP representing the cell distribution in the various subclusters (distinguishable by different colors) in WT and gata2b^+/−^ for HSPCs (A), Proliferative progenitors (B), Progenitors (C), Myeloid progenitors 1 (D), Myeloid progenitors 2 (E), Erythroid progenitors 2 (F), HSPCs and thrombocytes (G), and Erythroid progenitors 3 (H), with respective quantification of distribution between genotypes. Significantly differentially distributed subclusters in pink. FDR<0.05 & Log2 fold change >1.

## References

1. Gao X, Johnson KD, Chang YI, et al. Gata2 cis-element is required for hematopoietic stem cell generation in the mammalian embryo. J Exp Med. Dec 16 2013;210(13):2833–42. doi:jem.20130733 [pii] 20130733 [pii] 10.1084/jem.20130733

2. de Pater E, Kaimakis P, Vink CS, et al. Gata2 is required for HSC generation and survival. J Exp Med. Dec 16 2013;210(13):2843–50. doi:jem.20130751 [pii] 20130751 [pii] 10.1084/jem.20130751

3. Tsai FY, Keller G, Kuo FC, et al. An early haematopoietic defect in mice lacking the transcription factor GATA-2. Nature. Sep 15 1994;371(6494):221-6. doi:10.1038/371221a0

4. Donadieu J, Lamant M, Fieschi C, et al. Natural history of GATA2 deficiency in a survey of 79 French and Belgian patients. Haematologica. Aug 2018;103(8):1278–1287. doi:haematol.2017.181909 [pii] 1031278 [pii] 10.3324/haematol.2017.181909

5. Spinner MA, Sanchez LA, Hsu AP, et al. GATA2 deficiency: a protean disorder of hematopoiesis, lymphatics, and immunity. Blood. Feb 6 2014;123(6):809–21. doi:S0006-4971(20)36040-7 [pii] 2013/515528 [pii] 10.1182/blood-2013-07-515528

6. Ostergaard P, Simpson MA, Connell FC, et al. Mutations in GATA2 cause primary lymphedema associated with a predisposition to acute myeloid leukemia (Emberger syndrome). Nat Genet. Sep 4 2011;43(10):929–31. doi:ng.923 [pii] 10.1038/ng.923

7. Hsu AP, Sampaio EP, Khan J, et al. Mutations in GATA2 are associated with the autosomal dominant and sporadic monocytopenia and mycobacterial infection (MonoMAC) syndrome. Blood. Sep 8 2011;118(10):2653–5. doi:S0006-4971(20)40686-X [pii] 2011/356352 [pii] 10.1182/blood-2011-05-356352

8. Dickinson RE, Griffin H, Bigley V, et al. Exome sequencing identifies GATA-2 mutation as the cause of dendritic cell, monocyte, B and NK lymphoid deficiency. Blood. Sep 8 2011;118(10):2656–8. doi:S0006-4971(20)40687-1 [pii] 10.1182/blood-2011-06-360313

9. Hahn CN, Chong CE, Carmichael CL, et al. Heritable GATA2 mutations associated with familial myelodysplastic syndrome and acute myeloid leukemia. Nat Genet. Sep 4 2011;43(10):1012–7. doi:ng.913 [pii] 10.1038/ng.913

10. Mutsaers PG, van de Loosdrecht AA, Tawana K, Bodor C, Fitzgibbon J, Menko FH. Highly variable clinical manifestations in a large family with a novel GATA2 mutation. Leukemia. Nov 2013;27(11):2247–8. doi:leu2013105 [pii] 10.1038/leu.2013.105

11. Wang X, Muramatsu H, Okuno Y, et al. GATA2 and secondary mutations in familial myelodysplastic syndromes and pediatric myeloid malignancies. Haematologica. Oct 2015;100(10):e398–401. doi:haematol.2015.127092 [pii] 100e398 [pii] 10.3324/haematol.2015.127092

12. Rodrigues NP, Janzen V, Forkert R, et al. Haploinsufficiency of GATA-2 perturbs adult hematopoietic stem-cell homeostasis. Blood. Jul 15 2005;106(2):477–84. doi:S0006-4971(20)53269-2 [pii] 10.1182/blood-2004-08-2989

13. Ling KW, Ottersbach K, van Hamburg JP, et al. GATA-2 plays two functionally distinct roles during the ontogeny of hematopoietic stem cells. J Exp Med. Oct 4 2004;200(7):871–82. doi:jem.20031556 [pii] 20031556 [pii] 10.1084/jem.20031556

14. Butko E, Distel M, Pouget C, et al. Gata2b is a restricted early regulator of hemogenic endothelium in the zebrafish embryo. Development. Mar 15 2015;142(6):1050–61. doi:142/6/1050 [pii] DEV119180 [pii] 10.1242/dev.119180

15. Dobrzycki T, Mahony CB, Krecsmarik M, et al. Deletion of a conserved Gata2 enhancer impairs haemogenic endothelium programming and adult Zebrafish haematopoiesis. Commun Biol. Feb 13 2020;3(1):71. doi:10.1038/s42003-020-0798-3 [pii] 798 [pii] 10.1038/s42003-020-0798-3

16. Gioacchino E, Koyunlar C, Zink J, et al. Essential role for Gata2 in modulating lineage output from hematopoietic stem cells in zebrafish. Blood Adv. Jul 13 2021;5(13):2687–2700. doi:S2473-9529(21)00348-7 [pii] 10.1182/bloodadvances.2020002993

17. Avagyan S, Weber MC, Ma S, et al. Single-cell ATAC-seq reveals GATA2-dependent priming defect in myeloid and a maturation bottleneck in lymphoid lineages. Blood Adv. Jul 13 2021;5(13):2673–2686. doi:S2473-9529(21)00347-5 [pii] 2020/ADV2020002992 [pii] 10.1182/bloodadvances.2020002992

18. Ma D, Zhang J, Lin HF, Italiano J, Handin RI. The identification and characterization of zebrafish hematopoietic stem cells. Blood. Jul 14 2011;118(2):289–97. doi:S0006-4971(20)44778-0 [pii] 2010/327403 [pii] 10.1182/blood-2010-12-327403

19. Tamplin OJ, Durand EM, Carr LA, et al. Hematopoietic stem cell arrival triggers dynamic remodeling of the perivascular niche. Cell. Jan 15 2015;160(1-2):241–52. doi:S0092-8674(14)01638-9 [pii] 10.1016/j.cell.2014.12.032

20. Hao Y, Hao S, Andersen-Nissen E, et al. Integrated analysis of multimodal single-cell data. Cell. Jun 24 2021;184(13):3573–3587 e29. doi:S0092-8674(21)00583-3 [pii] 10.1016/j.cell.2021.04.048

21. Stuart T, Srivastava A, Madad S, Lareau CA, Satija R. Single-cell chromatin state analysis with Signac. Nat Methods. Nov 2021;18(11):1333–1341. doi:10.1038/s41592-021-01282-5 [pii] 10.1038/s41592-021-01282-5

22. Cao J, Spielmann M, Qiu X, et al. The single-cell transcriptional landscape of mammalian organogenesis. Nature. Feb 2019;566(7745):496-502. doi:10.1038/s41586-019-0969-x [pii] 10.1038/s41586-019-0969-x

23. Boatman S, Barrett F, Satishchandran S, Jing L, Shestopalov I, Zon LI. Assaying hematopoiesis using zebrafish. Blood Cells Mol Dis. Dec 2013;51(4):271–6. doi:S1079-9796(13)00160-5 [pii] 10.1016/j.bcmd.2013.07.009

24. Traver D, Paw BH, Poss KD, Penberthy WT, Lin S, Zon LI. Transplantation and in vivo imaging of multilineage engraftment in zebrafish bloodless mutants. Nat Immunol. Dec 2003;4(12):1238–46. doi:ni1007 [pii] 10.1038/ni1007

25. Langenau DM, Ferrando AA, Traver D, et al. In vivo tracking of T cell development, ablation, and engraftment in transgenic zebrafish. Proc Natl Acad Sci U S A. May 11 2004;101(19):7369–74. doi:0402248101 [pii] 1017369 [pii] 10.1073/pnas.0402248101

26. Renshaw SA, Loynes CA, Trushell DM, Elworthy S, Ingham PW, Whyte MK. A transgenic zebrafish model of neutrophilic inflammation. Blood. Dec 15 2006;108(13):3976–8. doi:S0006-4971(20)52189-7 [pii] 10.1182/blood-2006-05-024075

27. Ellett F, Pase L, Hayman JW, Andrianopoulos A, Lieschke GJ. mpeg1 promoter transgenes direct macrophage-lineage expression in zebrafish. Blood. Jan 27 2011;117(4):e49–56. doi:S0006-4971(20)58625-4 [pii] 2010/314120 [pii] 10.1182/blood-2010-10-314120

28. Page DM, Wittamer V, Bertrand JY, et al. An evolutionarily conserved program of B-cell development and activation in zebrafish. Blood. Aug 22 2013;122(8):e1–11. doi:S0006-4971(20)54254-7 [pii] 2012/471029 [pii] 10.1182/blood-2012-12-471029

29. Carmona SJ, Teichmann SA, Ferreira L, et al. Single-cell transcriptome analysis of fish immune cells provides insight into the evolution of vertebrate immune cell types. Genome Res. Mar 2017;27(3):451–461. doi:gr.207704.116 [pii] 10.1101/gr.207704.116

30. Danilova N, Bussmann J, Jekosch K, Steiner LA. The immunoglobulin heavy-chain locus in zebrafish: identification and expression of a previously unknown isotype, immunoglobulin Z. Nat Immunol. Mar 2005;6(3):295–302. doi:ni1166 [pii] 10.1038/ni1166

31. Kortum AN, Rodriguez-Nunez I, Yang J, et al. Differential expression and ligand binding indicate alternative functions for zebrafish polymeric immunoglobulin receptor (pIgR) and a family of pIgR-like (PIGRL) proteins. Immunogenetics. Apr 2014;66(4):267–79. doi:10.1007/s00251-014-0759-4

32. Macaulay IC, Svensson V, Labalette C, et al. Single-Cell RNA-Sequencing Reveals a Continuous Spectrum of Differentiation in Hematopoietic Cells. Cell Rep. Feb 2 2016;14(4):966–977. doi:S2211-1247(15)01538-7 [pii] 10.1016/j.celrep.2015.12.082

33. Tang Q, Iyer S, Lobbardi R, et al. Dissecting hematopoietic and renal cell heterogeneity in adult zebrafish at single-cell resolution using RNA sequencing. J Exp Med. Oct 2 2017;214(10):2875–2887. doi:jem.20170976 [pii] 20170976 [pii] 10.1084/jem.20170976

34. Athanasiadis EI, Botthof JG, Andres H, Ferreira L, Lio P, Cvejic A. Single-cell RNA-sequencing uncovers transcriptional states and fate decisions in haematopoiesis. Nat Commun. Dec 11 2017;8(1):2045. doi:10.1038/s41467-017-02305-6 [pii] 2305 [pii] 10.1038/s41467-017-02305-6

35. Wang J, Sun Q, Morita Y, et al. A differentiation checkpoint limits hematopoietic stem cell self-renewal in response to DNA damage. Cell. Mar 2 2012;148(5):1001–14. doi:S0092-8674(12)00145-6 [pii] 10.1016/j.cell.2012.01.040

36. Pauklin S, Vallier L. The cell-cycle state of stem cells determines cell fate propensity. Cell. Sep 26 2013;155(1):135–47. doi:S0092-8674(13)01025-8 [pii] 10.1016/j.cell.2013.08.031

37. Welch JJ, Watts JA, Vakoc CR, et al. Global regulation of erythroid gene expression by transcription factor GATA-1. Blood. Nov 15 2004;104(10):3136–47. doi:S0006-4971(20)55869-2 [pii] 10.1182/blood-2004-04-1603

38. Wang F, Zhu Y, Guo L, et al. A regulatory circuit comprising GATA1/2 switch and microRNA-27a/24 promotes erythropoiesis. Nucleic Acids Res. Jan 2014;42(1):442–57. doi:gkt848 [pii] 10.1093/nar/gkt848

39. Cantor AB, Orkin SH. Coregulation of GATA factors by the Friend of GATA (FOG) family of multitype zinc finger proteins. Semin Cell Dev Biol. Feb 2005;16(1):117–28. doi:S1084-9521(04)00102-8 [pii] 10.1016/j.semcdb.2004.10.006

40. Grass JA, Boyer ME, Pal S, Wu J, Weiss MJ, Bresnick EH. GATA-1-dependent transcriptional repression of GATA-2 via disruption of positive autoregulation and domain-wide chromatin remodeling. Proc Natl Acad Sci U S A. Jul 22 2003;100(15):8811–6. doi:1432147100 [pii] 1008811 [pii] 10.1073/pnas.1432147100

41. Pal S, Cantor AB, Johnson KD, et al. Coregulator-dependent facilitation of chromatin occupancy by GATA-1. Proc Natl Acad Sci U S A. Jan 27 2004;101(4):980–5. doi:0307612100 [pii] 1010980 [pii] 10.1073/pnas.0307612100

42. Bresnick EH, Lee HY, Fujiwara T, Johnson KD, Keles S. GATA switches as developmental drivers. J Biol Chem. Oct 8 2010;285(41):31087–93. doi:S0021-9258(19)88840-3 [pii] R110.159079 [pii] 10.1074/jbc.R110.159079

43. Monika Belickova M, Merkerova MD, Votavova H, et al. Up-regulation of ribosomal genes is associated with a poor response to azacitidine in myelodysplasia and related neoplasms. Int J Hematol. Nov 2016;104(5):566–573. doi:10.1007/s12185-016-2058-3 [pii] 10.1007/s12185-016-2058-3

44. Nakayama Y, Yamaguchi N. Multi-lobulation of the nucleus in prolonged S phase by nuclear expression of Chk tyrosine kinase. Exp Cell Res. Apr 1 2005;304(2):570–81. doi:S0014-4827(04)00703-7 [pii] 10.1016/j.yexcr.2004.11.027

45. Ullah Z, Lee CY, Depamphilis ML. Cip/Kip cyclin-dependent protein kinase inhibitors and the road to polyploidy. Cell Div. Jun 2 2009;4:10. doi:1747-1028-4-10 [pii] 10.1186/1747-1028-4-10

46. Haferlach T, Nagata Y, Grossmann V, et al. Landscape of genetic lesions in 944 patients with myelodysplastic syndromes. Leukemia. Feb 2014;28(2):241–7. doi:leu2013336 [pii] 10.1038/leu.2013.336

47. Rossi A, Kontarakis Z, Gerri C, et al. Genetic compensation induced by deleterious mutations but not gene knockdowns. Nature. Aug 13 2015;524(7564):230-3. doi:nature14580 [pii] 10.1038/nature14580

