## Supplementary material for "GATA2 haploinsufficiency causes an epigenetic feedback mechanism resulting in myeloid and erythroid dysplasi": Supplemantal fig 1-7

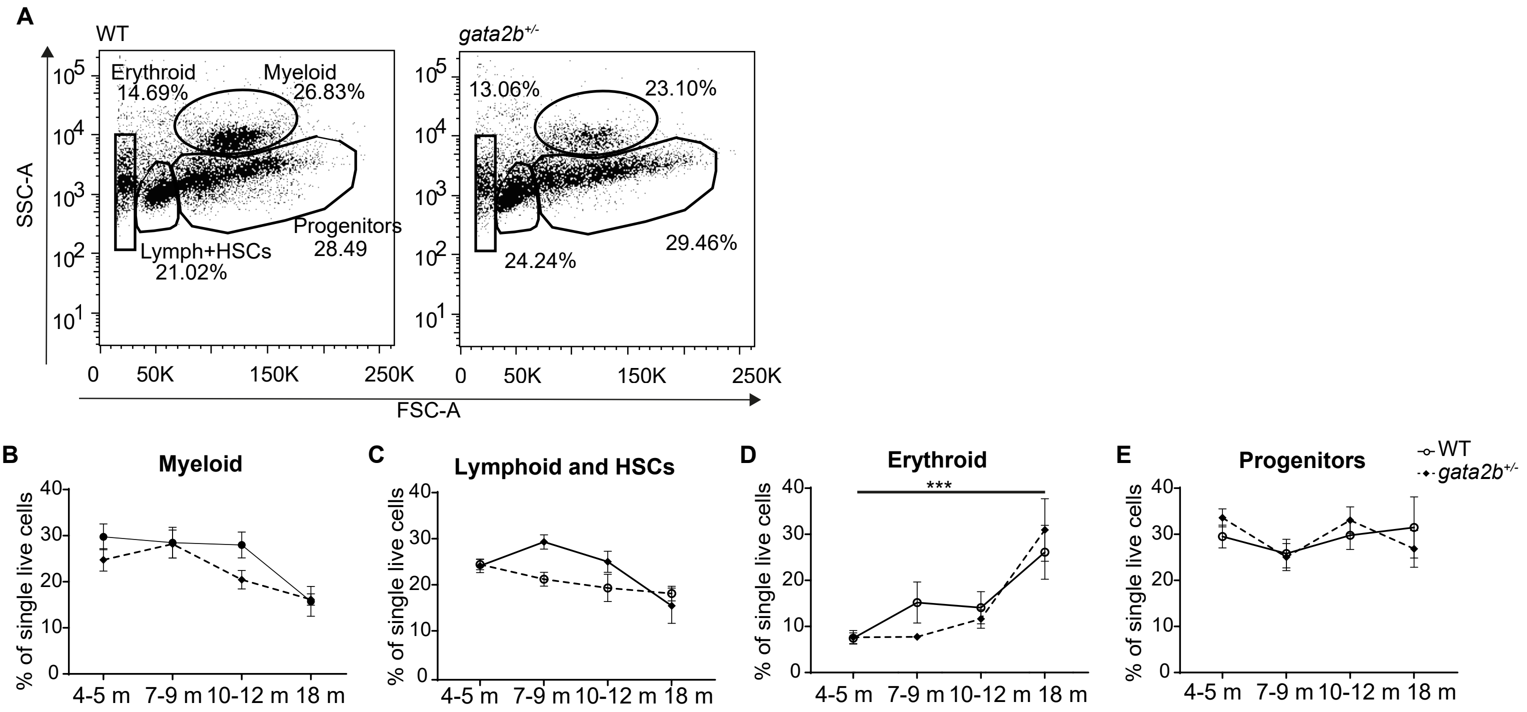


### Supplementary figure 1

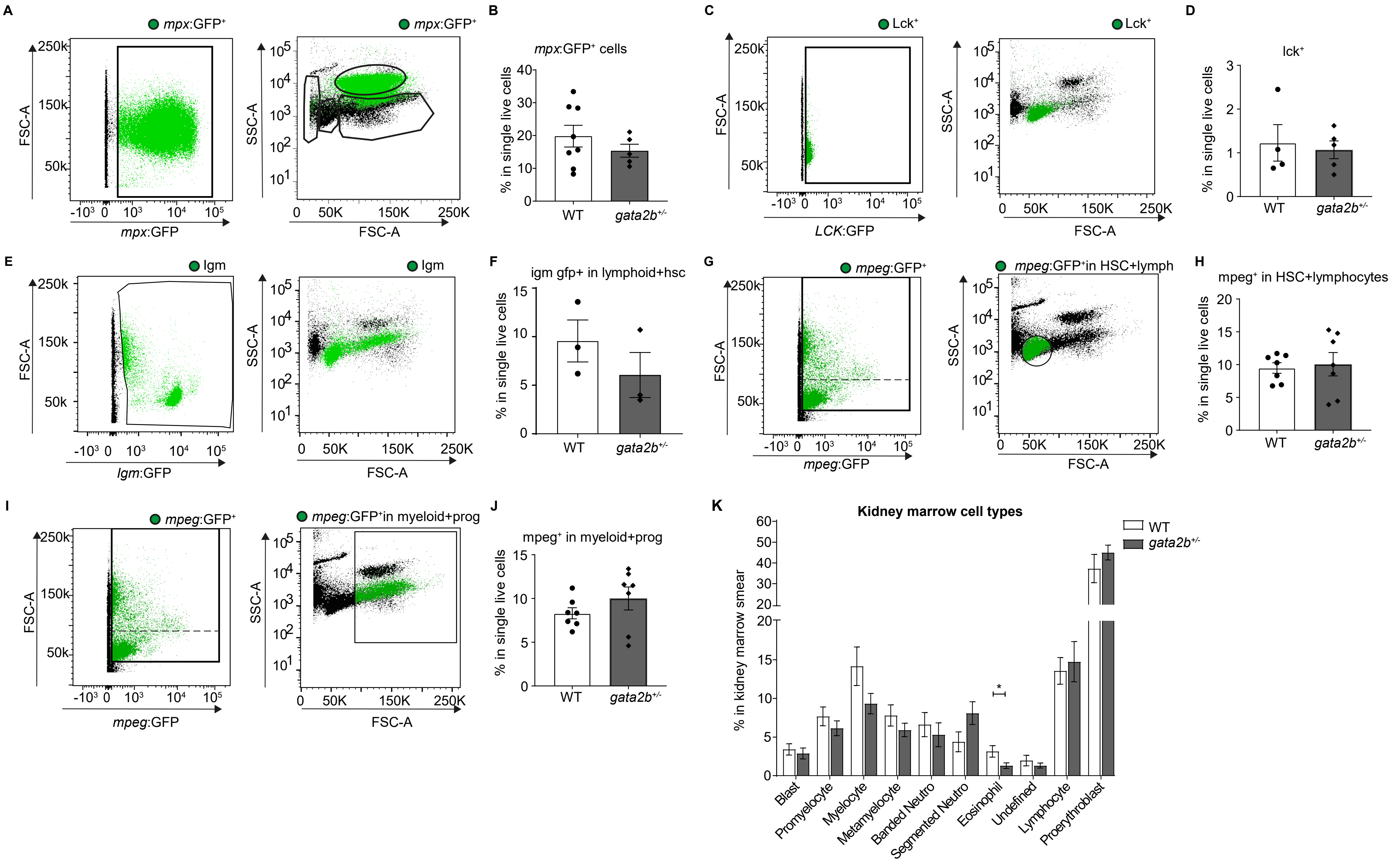


**Supplementary figure 2**

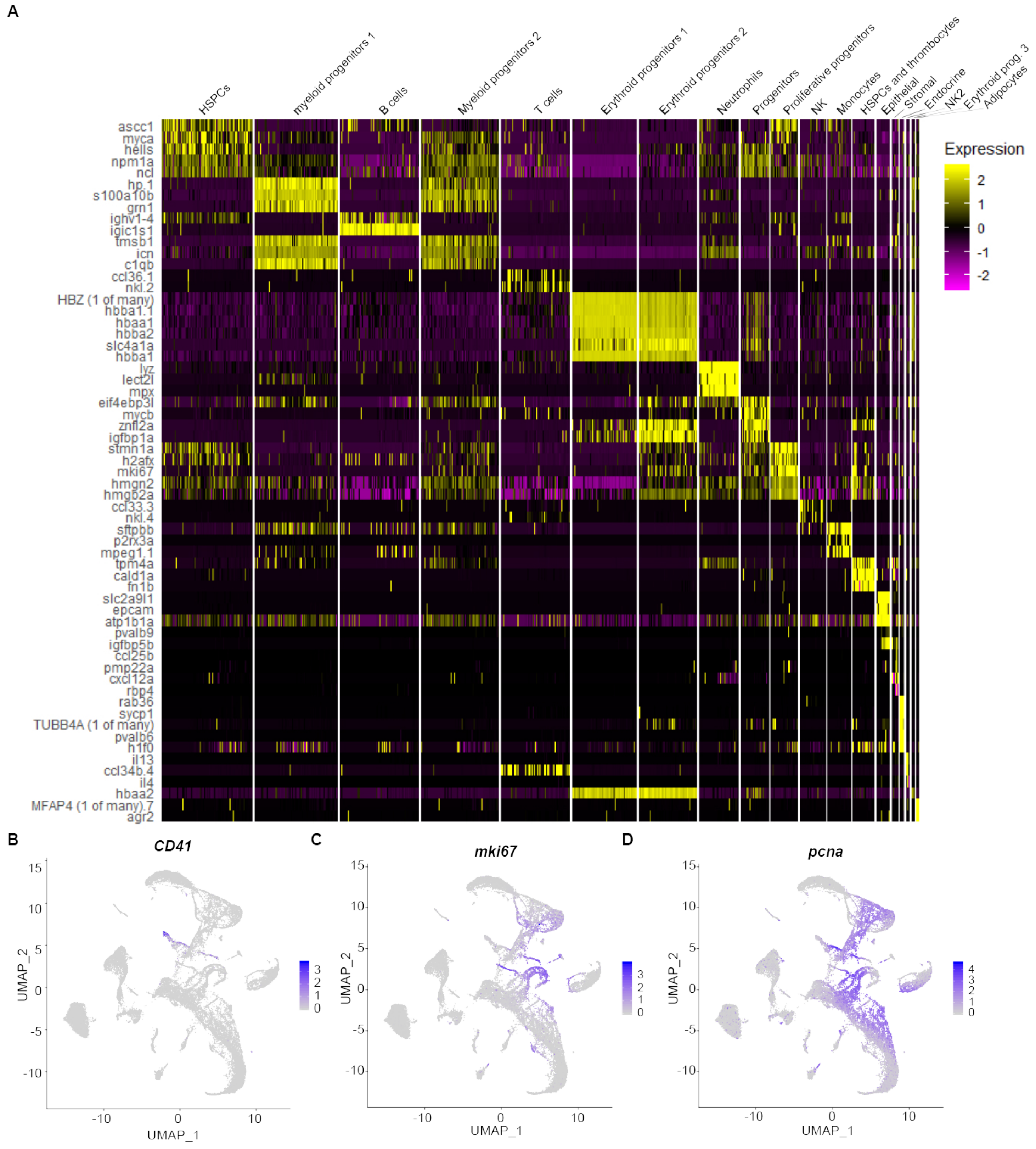


**Supplementary figure 3**

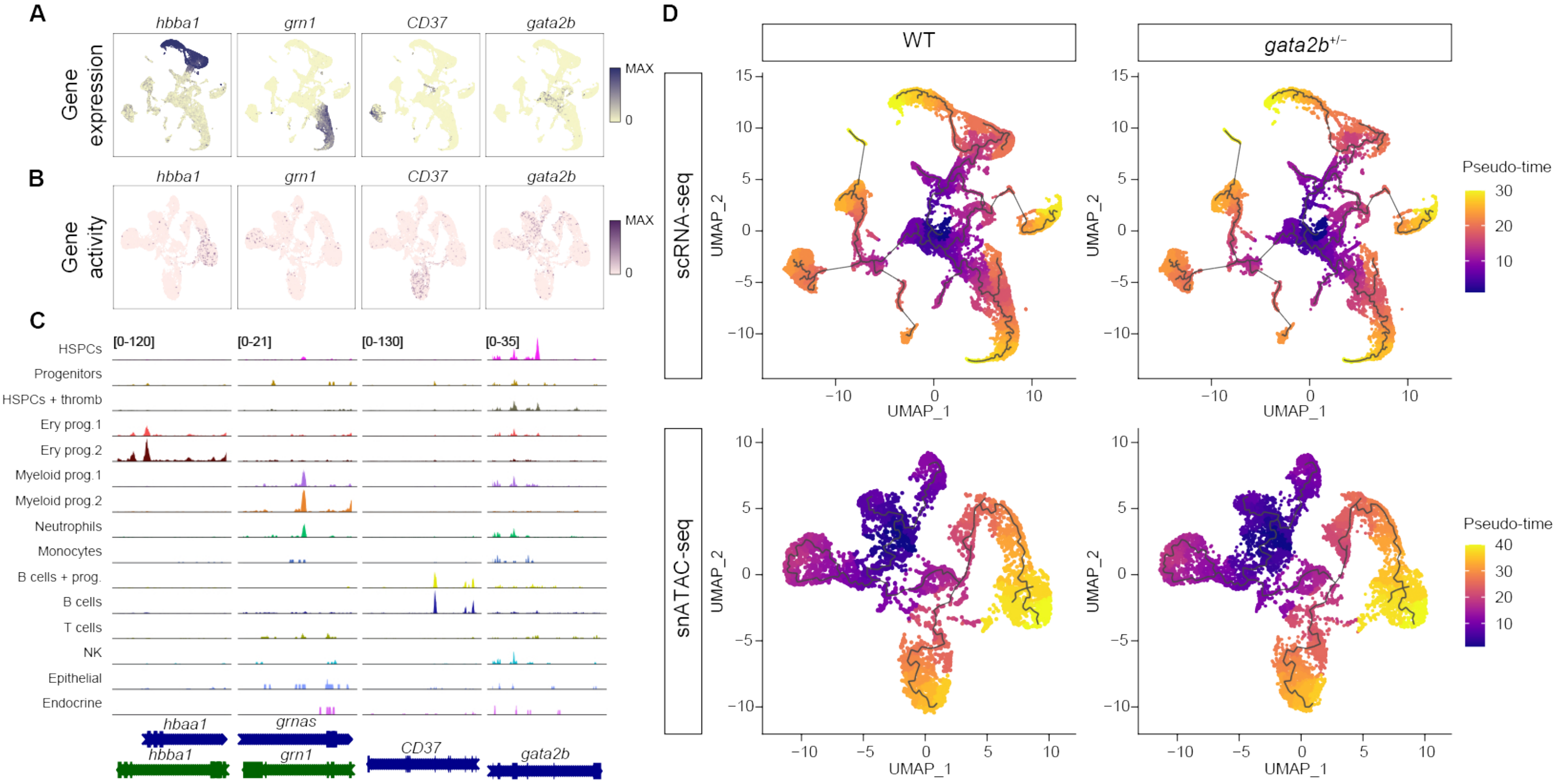


**Supplementary figure 4**

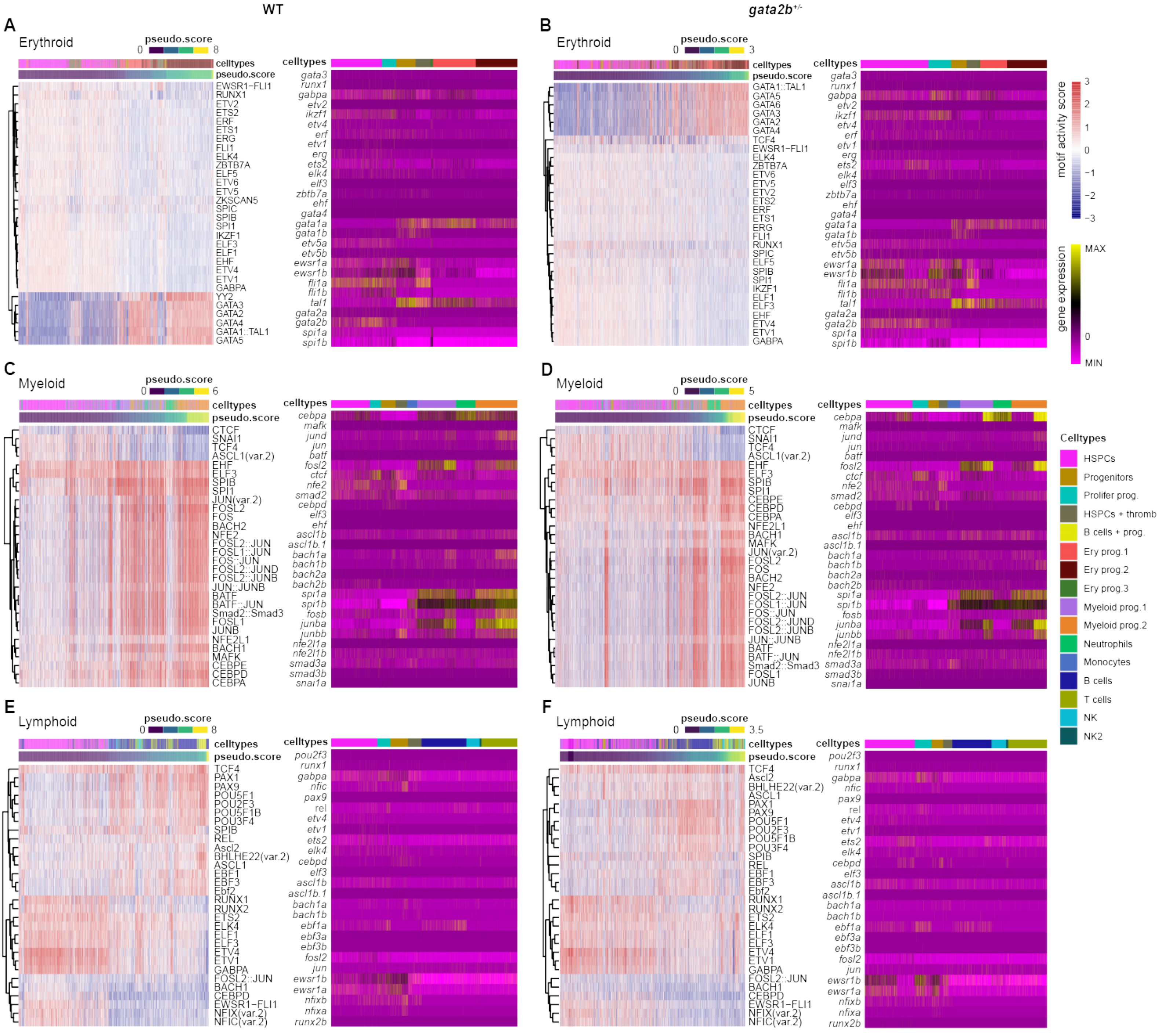


**Supplementary figure 5**

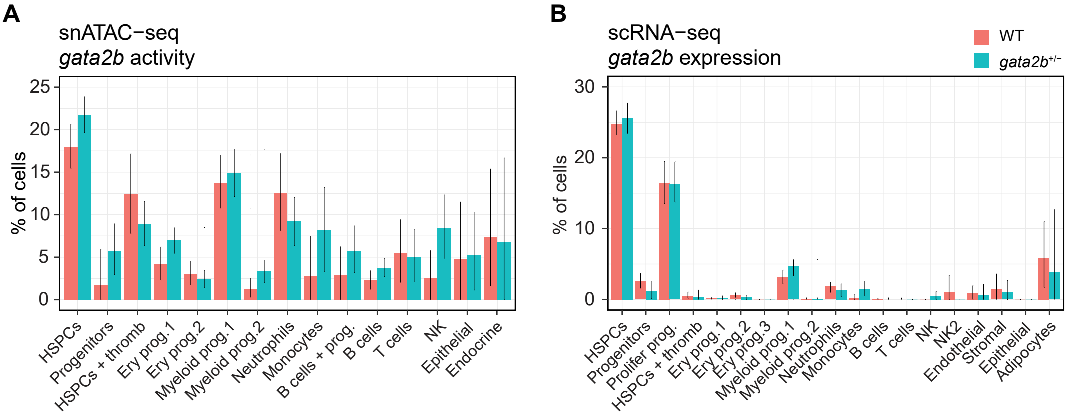


**Supplementary figure 6**

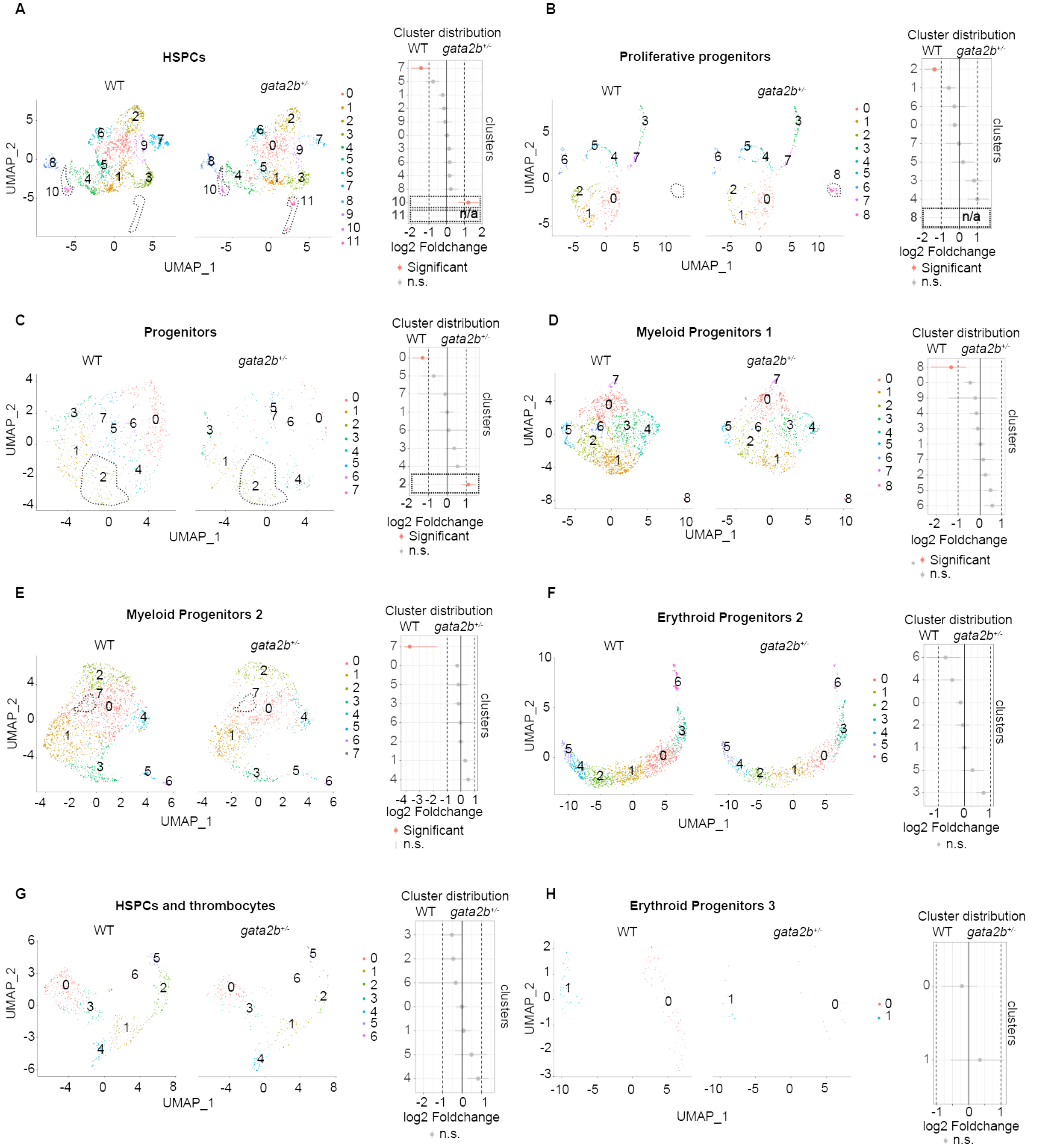


**Supplementary figure 7**
